# Aerobic scope does matter in temperature-size rule, but only under optimal conditions

**DOI:** 10.1101/2020.03.04.976050

**Authors:** Aleksandra Walczyńska, Anna Maria Labecka, Mateusz Sobczyk

## Abstract

We united the theoretical predictions on the factors responsible for the occurrence and evolutionary significance of the temperature-size rule. We tested the causal connection among them assuming that (i) the temperature-size rule is the response to temperature-dependent oxygenic conditions, (ii) body size decrease is a consequence of cell shrinkage in response to exposure to hypoxia, (iii) this response enables to keep the wide scope for aerobic performance, and (iv) it prevents the decrease in fitness. We conducted our tests on three clones of the rotifer *Lecane inermis* with different thermal preferences. These clones were exposed to three experimental regimes: mild hypoxia, severe hypoxia driven by a too high temperature, and severe hypoxia driven by an inadequate oxygen concentration. The results showed that our causative reasoning was generally correct, but only under mildly hypoxic conditions. In more stressful environments, rotifers had clone- and condition-specific responses, which in fact were equally successful in terms of the levels of fitness. Our results join for the first time all factors connecting the cause and effect in the temperature-size rule. They indicate the importance of the conditions under which it should be tested. The most important messages from this study were that (i) a decrease in the body size was one of but not the only option for preventing fitness reduction under hypoxia, and (ii) such a response to higher temperature enabled the maintenance of wide aerobic scope in clone-specific, thermally optimal conditions.

## Introduction

The phenotypic body size decrease with increasing temperature (temperature-size rule, TSR; Atkinson, 1994) has created a major puzzle for evolutionary ecologists (Berrigan and Charnov, 1994) because of its counterintuitive pattern. The reason for this confusion is that, according to the theory of optimal energy allocation, organisms growing in advantageous warm conditions should maximize their size to increase their own fitness, while those growing in harsh cold conditions should not postpone their maturity for long because of the risk of premature death. There have been different hypotheses on why the size response to temperature contradicts this pattern. Among them, the most promising is the hypothesis on the adaptive size response under oxygen deficiency (but see also Audzijonyte et al., 2019). The relative oxygen concentration decreases with increasing temperature (see Wetzel, 2001 for aquatic systems), and then the steeper increase in oxygen demands compared to oxygen supply causes oxygenic stress in ectothermic organisms (Atkinson et al., 2006; Verberk et al., 2011; Woods, 1999). The decrease in cell size has been suggested to increase the efficiency of oxygen transport to the mitochondria (Woods, 1999), and as a consequence, the body size shrinks as well. The adaptive hypothesis proposed was that temperature defines the window for the aerobic scope, namely the range of aerobic metabolism that can be performed under given conditions, and the ability to maintain the flexible efficiency of aerobic respiration prevents a possible decrease in fitness (Poertner, 2010). This mechanism was linked to the body size adjustment to temperature by Atkinson et al. (2006). The hypothesis on the oxygen-driven size-to-temperature response has been positively verified both indirectly (Berner et al., 2007; Harrison et al., 2010; Rollinson and Rowe, 2018a; Rollinson and Rowe, 2018b; Santilli and Rollinson, 2018; Verberk and Atkinson, 2013) and directly (Czarnoleski et al., 2015; Frazier et al., 2001; Hoefnagel and Verberk, 2015). However, results relating this pattern directly to organismal fitness are scarce (Prokosch et al., 2019; Walczyńska et al., 2015a), while such a reference is the only way that enables reliable conclusions on the possible evolutionary meaning of TSR.

In this study, we linked the hypothesized proximate and ultimate mechanisms behind the phenotypically driven size decrease with increasing temperature for the case of the rotifer *Lecane inermis*. We aimed to thoroughly test the theoretical reasoning behind hypoxia driving body size shrinkage to prevent the reduction in fitness, which is presented in the literature. Because the TSR has been empirically confirmed to be condition-sensitive and to be followed only in an optimal thermal range (Walczyńska et al., 2016), we conducted our tests under optimal and suboptimal conditions; the latter was further divided between hypoxia driven by stress-inducing high temperatures or by the stress-inducing low oxygen levels. This distinction between mild *vs*. harsh conditions enabled inference on the possible thresholds for TSR performance, additionally referred to fitness.

We examined the aerobic scope, our hypothesized ultimate mechanism, by estimating the specific dynamic action (SDA). This measure refers to the increasing metabolic expenditures in animals and is associated with ingestion, digestion, assimilation and absorption of food (McCue, 2006; Secor, 2009), when the increasing oxygen consumption was mostly associated with the biochemical transformation of food and the synthesis of the new proteins (Jordan and Steffensen, 2007). In practice, it was estimated as a difference in the respiration rate between the fed and hungry individuals that cannot be maintained at rest. The hypothesized link between aerobic scope and SDA is that the larger the oxygen consumption associated with SDA, the wider the capacity of aerobic metabolism (aerobic scope). This assumption was previously used for the case of *Daphnia magna* (Chopelet et al., 2008). Our approach was to compare the oxygen consumption rate in hypoxia for animals previously exposed to normoxic or hypoxic conditions. We expected that hypoxia would lower the SDA amplitude in hypoxia-naïve animals compared to hypoxia-experienced animals. This hypothesis was based on Fry (1947), who wrote that dissolved oxygen acts as a limiting factor for metabolic rate; thus, its reduction decreases aerobic scope and, as a consequence, the energy budgets of aquatic breathers. Such an arresting effect of hypoxia on SDA width was previously reported for a cod fish (Jordan and Steffensen, 2007) and carabid beetles (Gudowska and Bauchinger, 2018). We hypothesized that hypoxia selects for small cell size, followed by small body size, to maintain high effectiveness of aerobic metabolism and to maximize fitness (Fig. 1A). We expected this pattern to differ across the hypoxia treatments, with the more direct response under mild hypoxia. Therefore, our main goal was to compare all traits examined in rotifers exposed to normoxia and those exposed to hypoxia, across three experimental treatments.

**Fig. 1.**
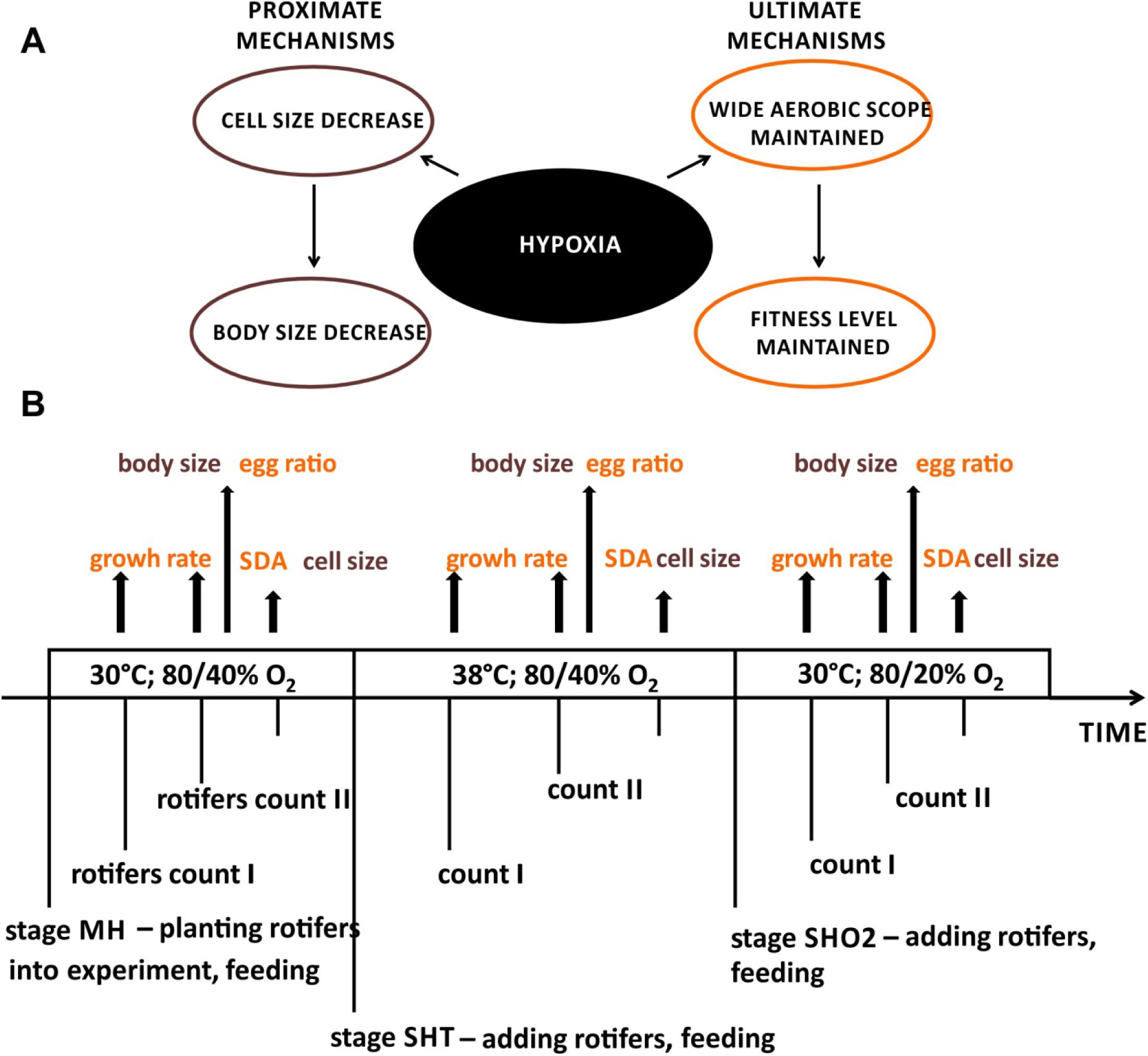
Concept of the study. A – hypothetical effect of hypoxia on the traits estimated: hypoxia causes cell size and body size decrease (proximate mechanisms) to maintain the wide aerobic scope to prevent the possible fitness decline (ultimate mechanisms). B – Experimental setup: the study was divided into three subsequent stages with different hypoxic conditions; five traits were investigated within three days in each stage. SDA – specific dynamic action (a proxy for aerobic scope); MH stage – hypoxia type = mild hypoxia; SHT stage – hypoxia type = severe hypoxia with high temperature; SHO2 stage – hypoxia type = severe hypoxia with low oxygen concentration.

Apart from conducting the study in optimal or stressful conditions, we also tested for possible differences in the responses of animals acclimated to different thermal conditions: cold, intermediate, and warm. Such a distinction addresses an issue of the possible linkage between the thermal preferences and thermally induced body size adjustment. We used three clones of the rotifer *L. inermis*, a species that has previously been found to follow the TSR in the laboratory and in the field (Kiełbasa et al., 2014), to show the adaptive body size response to thermally driven oxygen conditions based on the cell size adjustment (Walczyńska et al., 2015a) and to perform simple two-point TSR control within its lifecycle, by the mother and during egg development (Walczyńska et al., 2015b). To our knowledge, this is the first attempt to test for the proximate and ultimate mechanisms behind the TSR in such a comprehensive way.

## Materials and Methods

### Clones studied

The *Lecane inermis* Bryce rotifer species has previously been tested for different aspects of size response to thermal and oxygenic conditions. Its thermal performance curve was also examined, and the optimal temperature was estimated to be 30°C (Walczyńska et al., 2016). This species is very convenient for the studies on the changes in phenotypic response between the maternal and offspring generations because of its short generation time, estimated previously for the range between 15 °C and 25 °C as two days (Walczyńska et al., 2015b). We chose three *L. inermis* clones that were founded from one individual each, which were isolated from activated sludge from a small wastewater treatment plants in southern Poland. The clones differed in their acclimation conditions due to exposure to laboratory conditions representing relatively low, intermediate or high temperatures. Thus, we assumed the existence of differences in thermal preferences among the clones. Our predictions were confirmed in a previous study conducted for six clones (Stuczyńska et al., 2020). A general pattern found was the linkage between the pre-experimental thermal exposure and clonal thermal preference during the study. The three clones used here refer to clones Cold, Int2 and Warm2 from the previous study and are here named Cold for the cold preference, Int for intermediate temperature preference and Warm for high heat preference, respectively. Preceding the experiment, all clones were kept in Petri dishes at 25°C in the dark in Żywiec spring water (Poland) and fed patented Novo powder (a patent by Pajdak-Stós et al., 2017).

### Experimental setup

The experiment was divided into three parts that constituted different hypoxic regimes (Fig. 1 B): MH – mild hypoxia (30°C and 80% or 40% O_2_) was followed by SHT – severe hypoxia with temperature above the optimum (38°C and 80% or 40% O_2_) and then by SHO2 – severe hypoxia with very low oxygen level (30°C and 80% or 20% O_2_). In each part, the three rotifer clones were exposed to either normoxia (80% O_2_) or hypoxia (40% or 20% O_2_). The choice of 80% normoxia instead of 100% was dictated by the fact that it is much easier to maintain stable 80% than 100% oxygen concentrations, and in this case, we were assured that the rotifers were not exposed to fluctuating oxygen conditions that could act as an uncontrolled factor. The relative 80% oxygen concentration was equivalent to approximately 6 mg/L O_2_, which is very close to the optimal value for this species living in nature (7.5 mg/L O_2_) (Berzins and Pejler, 1989) and much higher than the oxygen concentration frequently set under the activated sludge conditions (2 mg/L O_2_)(Kiełbasa et al., 2014) from which all the studied clones were established. The value of 2-3 mg/L O_2_ was also provided as a standard threshold for hypoxia (reviewed in Hrycik et al., 2017).

The experimental conditions were set in four temperature-regulated water baths (Memmert, Germany) connected to a 4-channel OxyReg O_2_ regulation system (Loligo Systems, Denmark) calibrated to 0% and 100% air-saturated water. A stable level of hypoxia was maintained by adding pure (comestible) nitrogen to the baths. The relative oxygen concentrations of 80, 40 and 20% O_2_ were equivalent to approximately 17% (∼6.0 mg/L), 8% (∼2.8 mg/L) and 4% (∼1.5 mg/L) of the atmospheric oxygen, respectively. Two baths were set to normoxia and two to hypoxia, with the same temperature in all. Six 100 mL cages made of 5 µm mesh enclosed in slide frames sealed with silicone were placed in each water bath (two cages-replicates per clone in each bath). This type of cage has been successfully used previously (Walczyńska et al., 2015a).

Each water bath was filled with Żywiec spring water as the medium. The initial densities of rotifers planted at the beginning of experimental parts were 110, 120 and 250 indiv./mL for MH, SHT and SHO2 stages, respectively. These numbers were dictated by the clone that was the least numerous at the moment of counting; our aim was to start with the highest possible number of rotifers, even across clones. Such initial densities of rotifers were far below the population carrying capacity because *L. inermis* rotifers may reach densities as high as tens of thousands per mL (45 000 indiv/mL was reported by Walczyńska et al., 2017). The rotifers in each cage were fed 200 µL of commercial bioproduct Bio-Trakt®, which has been successfully used in *L. inermis* mass culture previously (Fiałkowska et al., 2019). During or just after the three days of each experimental stage, we took samples to estimate the physiological (specific dynamic action (SDA), which is a proxy for aerobic scope, and nucleus size, which is a proxy for cell size, and life history traits (body size, population growth rate, egg ratio), as presented in Fig. 1B.

### Oxygen consumption

The hypothesis of the decrease in size under hypoxic conditions to maintain a wide aerobic scope was tested by estimating the SDA. This physiological difference in respiration rate between fed and hungry animals was previously used as a proxy for aerobic scope in the case of *Dapnia magna* (Chopelet et al., 2008). We estimated the SDA using spectrophotometry (Orion™ AquaMate 8000 UV-VIS, Thermo Scientific, USA) with a modified Winkler titration (Carpenter, 1965; Chopelet et al., 2008; Roland et al., 1999), which has been utilized as a highly accurate method for determining dissolved oxygen (Helm et al., 2012). In this method, the absorbance measure is proportional to the amount of oxygen that is dissolved in a sample as a consequence of the chemical reaction chain. For each experimental stage, we sampled the rotifers from the normoxia and hypoxia treatments and measured the aerobic scope of both groups exposed to hypoxic conditions, respective to the given stage. We expected that the rotifers exposed to experimental hypoxia in the preceding three days should have wider SDAs in the test under hypoxia than the normoxia-exposed rotifers. Two clones, Cold and Warm, were examined for SDA because the Int clone did not proliferate well enough to achieve reasonable sample sizes. On the third day of each experimental stage, we collected 10 mL samples from each cage and pooled the two replicates within the clone×oxic regime combination. For logistical reasons, the clones were analysed separately but within the same day for the cases of SHT stage and SHO2 stage. Because we did not conduct an analysis for both clones after the MH stage, we repeated this stage at the end of the experiment to collect the missing data on SDA for the Cold clone.

Each clone was divided into two 50 mL crystallizers with a 5 mL rotifer sample and 15 mL of hypoxic medium taken directly from the respective water bath representing the experimental hypoxic conditions. By applying this procedure, we diluted the nutritional particles sampled along with the rotifers. The rotifers in one crystallizer were then fed with 10 µL of Bio-Trakt® per 1 mL of sample (treatment: fed). The rotifers in the second crystallizer were not fed (treatment: hungry). The random division of clone samples into the fed and hungry treatments caused that the average rotifer body size was similar in both regimes. Therefore, we could assume that the possible differences in oxygen consumption were not related to size differences.

Both samples were incubated for 2 h at the same temperature the rotifers experienced during the respective experimental stage. The relatively short exposure to starving conditions was dictated by the fact that rotifers become starved relatively quickly and that severe starvation has a very profound effect on metabolism (reviewed by Galkovskaya, 1995).

After this period, from both treatments, we fished the swimming, healthy-looking rotifers in bunches of 30 and transferred them to the wells of 24-well tissue plates filled with membrane made of parafilm, with four or five replicates (= rotifer bunches) per treatment. The rotifers from each well were then quantitatively transferred to individual 2 mL autosampler vials (Agilent Technologies, USA) with hypoxic medium taken directly from one experimental water bath with hypoxic conditions. The vials (five replicates per fed and hungry treatments) were filled according to the following procedure: 1 mL of medium, the rotifer sample from a well, and another 1 mL of medium until a convex meniscus was achieved. The vials were then closed with dedicated caps punctured beforehand with a sterile needle to remove excess fluid and air. The control tubes were prepared following the same procedure but without the animals. The samples were incubated for another two (38°C) or three (30°C) hours in hypoxic conditions, with the level of hypoxia set to 40%, 40% and 20% O_2_ for the experimental MH, SHT and SHO2 stages, respectively.

After incubation, the Winkler chemicals were added following the procedure adopted from Carpenter (1965): 16 µL of MnCl_2_·4H_2_O (3 M), 16 µL of NaI (4 M) in NaOH (8 M), and, after vigorous shaking to cause precipitation, 16 µL of H_2_SO_4_ (5 M). The vials were then shaken to dissolve the precipitate, and 800 µL subsamples were taken directly to spectrophotometric micro-cuvettes (Brand, Germany). The absorbance was measured immediately at a wavelength of 440 nm within the first 10 minutes to prevent iodine evaporation (Roland et al., 1999). The calibration curve for absorbance was estimated for 0, 15, 35 and 50% O_2_ (in relative values), with 5 replicates per oxygen level. In the final estimation, the extreme values were excluded, and the average was calculated for the remaining three values per oxygen level. The calibration curve fitting was estimated at R^2^ = 0.9953.

The control samples were checked for their reliability for each combination of hypoxia type and clone separately. In all but two cases, the controls showed similar and reliable patterns. In the case of the MH treatment with the Warm clone, the control samples showed a pattern indicating some sort of methodological error (supplementary materials, Fig. S1), and only one value attained for the first sample was taken into consideration. In the case of the SHT treatment with the Cold clone, one control sample had a very low value (Fig. S1), which made it unreliable. The final control value was calculated as a mean from the remaining two samples. SDA was estimated both as a difference in raw absorbance gained for the fed rotifers and that for hungry rotifers (absolute SDA) and as their division (factorial SDA). The comparison of these two approaches for the aerobic scope calculation was suggested by Clark et al. (2013).

The amount of consumed O_2_ was calculated from % to mg/L using the equation describing the relationship between these two variables provided by the OxyReg system. This was possible only for 30°C, due to the relatively wide range of O_2_ (%) covered (from approximately 19 to 95%), while the much shorter range for 38°C made this estimation less reliable.

### Cell size

We measured the nucleus size, a proxy for cell size, according to the method developed and used before for the same species (Walczyńska et al., 2015a). Such a procedure was dictate by the fact that cell size was reported to correlate with DNA size (Jalal et al., 2013; Kozlowski et al., 2003), meaning that nucleus size might be a convenient estimate of cell size (reviewed by Cavalier-Smith, 2005). The rotifers were sampled on the third day of each experimental part. Approximately 50 swimming rotifers were fished and placed in 1 mL Eppendorf tubes, and the samples were fixed with 10% buffered neutral formalin (Avantor Performance Materials Poland S.A., Poland). The staining procedure was based on nuclear red (Carl Roth, Germany), as described in Walczyńska et al. (2015a). We measured two perpendicular diameters of 10 nuclei per rotifer using ImageJ (NIH, USA), and we calculated their product. The nuclei represent different animal organs. In this context, it is worth noting that rotifers do not possess any specialized respiratory cells, and they respire through the whole body surface (Galkovskaya, 1995).

### Body size

We estimated the strength of the response to hypoxia in comparison to normoxia by measuring the body size. For this purpose, we sampled the rotifers on the second day of the experiment and fixed them with Lugol solution. The rotifers were photographed (Eclipse 80i microscope (Nikon, Japan) equipped with a DS-Ri2 camera assisted by NIS-Elements D software (Nikon)). The length and width were measured in ImageJ, and their product acted as a proxy for body area, as previously applied to the rotifers *Brachionus* sp. (Walczyńska and Serra, 2014) and *L. inermis* (Walczyńska et al., 2015a).

### Fitness measures

Two measures of fitness were compared: population growth rate and egg ratio, which is the number of eggs laid by an average female. To estimate the former, we sampled 1 mL from each cage (two cages × two water baths gave four replicates per clone×oxic regime combination), fixed with Lugol solution and counted the rotifers. These procedures were used twice: one day (count I) and after two days (count II) after the onset of the stage. During count II, the egg number was noted along with the female number for egg ratio estimation.

### Statistical analyses

All analyses were conducted in SAS 9 (SAS, 2013) in PROC MIXED, method = reml. In the case of SDA, the GLM model included four fixed factors: hypoxia type (experimental stage), oxic regime of rotifer origin (normoxia *vs*. hypoxia), and clone and food regime (fed *vs*. hungry), along with all their interactions. The non-significant interactions in the model were excluded using backward elimination.

Possible differences in the nuclei size and body size were tested separately in GLM models that included three fixed factors: hypoxia type, clone and oxic regime, and their interactions. The non-significant interactions were excluded using backward elimination. In the case of body size, the differences in size among the hypoxia types caused the considerable lack of homogeneity of variance, which prevented statistical analysis. Therefore, the data were standardized by recalculating them to the units of standard deviation for each experimental stage (= hypoxia type) separately.

Both fitness measures were analysed with ANCOVA (PROC MIXED) and were not analysed as ratios, which has been cited as a more appropriate method (Raubenheimer, 1995; Raubenheimer and Simpson, 1992). The models included three fixed factors: hypoxia type, clone and oxic regime, and their interactions. In the case of population growth rate *r*, the dependent variable was the rotifer number from count II, while that from count I was a covariate. In the case of the egg ratio, the dependent variable was the number of eggs, while a covariate was the number of rotifer females in count II. The dependent variables and covariates were natural logarithm-transformed. In the initial stages, we included all the possible first-level interactions of the covariates with the main factors to test whether they were appropriately used in the models. In both models, the covariates were found to significantly affect the dependent variables and to not interact with the main factors. In the next step, we constructed the models of all the main factors and possible interactions, plus a covariate. The non-significant interactions were removed using backward elimination.

## Results

### Oxygen consumption

The results for the absolute oxygen consumption, which was calculated from the equation O_2_ [mg/L] = 0.0752 × O_2_ [%], were compared with other published data for different rotifer species obtained from the literature (Table 1). Generally, the oxygen consumption rate for the units provided in Table 1 was approximately 4 times higher in fed rotifers than in hungry rotifers. This estimated difference was previously shown to be 2-3-fold (Galkovskaya, 1995), although the units and starvation times for this comparison were unclear. The oxygen consumption differed among hypoxia types, clones, rotifer origins (normoxia or hypoxia), and food regimes, as well as for some interactions (Table 2). Most interestingly, the O_2_ consumption of the hungry and fed rotifers significantly differed between the clones and the hypoxia types, according to the significant *hypoxia type×clone×food* interaction (Table 2). In each case, the fed rotifers consumed more oxygen than the hungry rotifers due to the metabolic costs associated with SDA. The ceiling of these costs was relatively constant across hypoxia types in the case of the Warm clone and was elevated in cold rotifers exposed to hypoxia driven by stress-inducing high temperatures. The SDA-related costs were generally higher in the rotifers that had experienced hypoxia in the experiment than in hypoxia-naïve rotifers (Fig. 2A). The negative value for the Warm clone from normoxia in the MH treatment was caused by two elevated points (see supplementary materials, Fig. S1), but the analysis for outliers showed that there was no objective reason for their removal from the dataset. The pattern for the SDA was similar regardless of the measure (whether absolute or factorial; Fig. 2). Generally, the width of the SDA across the hypoxic treatments was similar for the Cold clone, while in the Warm clone, the values were much higher under mild hypoxia than under the two severe hypoxia treatments. In mildly hypoxic conditions, both clones showed wider SDAs when previously exposed to hypoxia during the experiment than those that were hypoxia-naïve. This pattern was not repeated in the following severe hypoxia treatment. The cold clone maintained relatively high and similar SDA, regardless of previous hypoxia experience, while the warm clone displayed a clearly narrower SDA in the rotifers experiencing hypoxia than in those taken from the normoxia treatment (Fig. 2).

**Table 1.** Comparison of the absolute oxygen consumption measured under different conditions in different rotifer species.

| Rotifer species | O<sub>2</sub> cons.<br>[mg×indiv<sup>-1</sup>×h<sup>-1</sup>] | O<sub>2</sub> cons.<br>[mL×indiv<sup>-1</sup>×h<sup>-1</sup>] | Measurements<br>conditions | References |
| --- | --- | --- | --- | --- |
| *Asplanchna priodonta* |  | 0.48 × 10<sup>-5</sup> and<br>0.71 × 10<sup>-5</sup> | 20°C, well-fed rotifers<br>and after 28 h<br>starvation, respectively | Kirk et al. (1999) |
| *Asplanchna silvestrii* |  | 1.65 × 10<sup>-5</sup> and<br>1.54 × 10<sup>-5</sup> | 20°C, well-fed rotifers<br>and after 57 h<br>starvation, respectively | Kirk et al. (1999) |
| *Brachionus calyciflorus* |  | 0.16-0.62 × 10<sup>-5</sup> | Large numbers,<br>unknown age<br>distribution, different<br>conditions | Pourriot and<br>Deluzarches (1970),<br>after Doohan (1973) |
| *Brachionus calyciflorus* |  | 0.22 × 10<sup>-5</sup> | - | Belyatskaya (1959),<br>after Doohan (1973) |
| *Brachionus calyciflorus* |  | 0.5 × 10<sup>-5</sup> | - | Galkovskaya (1963)<br>after Doohan (1973) |
| *Brachionus calyciflorus* |  | 0.31 × 10<sup>-5</sup> and<br>0.13 × 10<sup>-5</sup> | 20°C, well-fed rotifers<br>and after 53 h<br>starvation, respectively | Kirk et al. (1999) |
| *Brachionus plicatilis* |  | 0.27 × 10<sup>-5</sup> | female without eggs;<br>20°C, sea water with no<br>food | Doohan (1973) |
| *Brachionus plicatilis* |  | 0.1-0.7 × 10<sup>-5</sup> | 20‰ salinity,<br>20°C | Hirata and Yamasaki<br>(1987) |
| *Keratella quadrata* | 0.08 × 10<sup>-5</sup> and<br>0.11 × 10<sup>-5</sup> |  | 14.5-18°C and 20°C,<br>respectively | Vinberg (1937), after<br>Galkovskaya (1995) |
| *Lecane inermis*, two<br>clones | 0.09 × 10<sup>-5</sup> and<br>0.38 × 10<sup>-5</sup> | 0.07 × 10<sup>-5</sup> and<br>0.29 × 10<sup>-5</sup> | 30°C, 2 h starved and<br>fed, respectively | this study |
| *Polyarthra dolichoptera* |  | 0.09 × 10<sup>-5</sup> and<br>0.027 × 10<sup>-5</sup> | 2 and 5°C, respectively | Galkovskaya (1995) |
| *Synchaeta pectinata* |  | 0.21 × 10<sup>-5</sup> and<br>0.22 × 10<sup>-5</sup> | 20°C, well-fed rotifers<br>and after 18 h<br>starvation, respectively | Kirk et al. (1999) |

**Table 2.** Final results of the GLM model for oxygen consumption in two *L. inermis* clones.

| Effect | F-value (df num, df den) | P-value |
| --- | --- | --- |
| hypoxia type | (2, 91) 38.81 | < 0.0001 |
| clone | (1, 91) 7.69 | 0.0067 |
| oxic regime | (1, 91) 100.88 | < 0.0001 |
| food regime | (1, 91) 643.58 | < 0.0001 |
| h. type × clone | (2, 91) 20.15 | < 0.0001 |
| h. type × oxic regime | (2, 91) 0.63 | 0.5371 |
| clone × oxic regime | (1, 91) 0.81 | 0.3698 |
| h. type × food regime | (2, 91) 4.81 | 0.0104 |
| clone × food regime | (1, 91) 2.59 | 0.1108 |
| oxic regime × food regime | (1, 91) 0.01 | 0.9034 |
| h. type × clone × food regime | (2, 91) 8.21 | 0.0005 |
| h. type × oxic regime × food regime | (2, 91) 3.88 | 0.0240 |

hypoxia type – subsequent experimental stages, standardized within each stage to the SD units oxic regime – the origin of the animals (rotifers could originate from the normoxic or hypoxic experimental conditions, but their respiration was always measured under hypoxia) food regime – fed or hungry rotifers

**Fig. 2.**
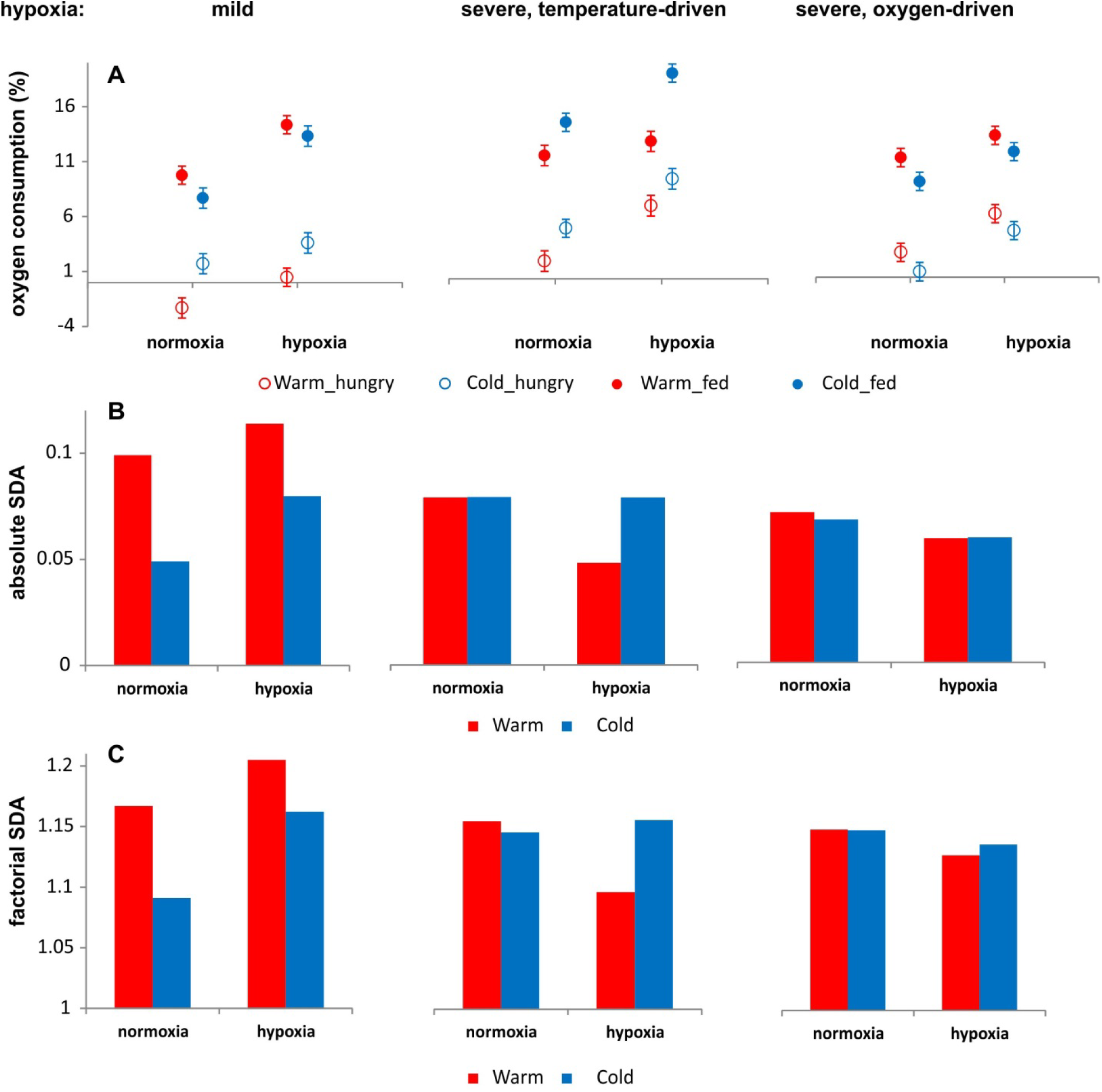
Results for oxygen consumption (in %), absolute aerobic scope and factorial aerobic scope for warm and cold clones across three regimes of different hypoxia types. A – oxygen consumption for rotifers exposed to experimental hypoxia or normoxia, then divided into groups of fed or hungry and tested for respiration rates under hypoxic conditions (4 or 5 replicates per group tested). Least square means ± s.e.. B – absolute scope for aerobic metabolism as a difference between the oxygen consumption of fed and hungry rotifers; absolute SDA. C – factorial scope for aerobic metabolism as a division of the amount of oxygen consumed by the fed rotifers to that consumed by the hungry rotifers, factorial SDA.

### Cell size

In total, we measured 1386 nuclei in 143 animals. For the experimental SHO2 stage, we were only able to take measurements for the Warm clone; the samples of other two clones were lost, most probably because of some unknown error with fixing. The mean nuclei size of 1.99 µm^2^ ± 0.62 SD was close to the 1.62 µm^2^ achieved previously at 32°C for the different *L. inermis* clones (Walczyńska et al., 2015a). We found a significant effect of the type of hypoxia and its interaction with the clone and oxygenic conditions on the nuclei size (Table 3), yet there was no effect of the oxygenic conditions alone. The clones differed in cell size, and their diverse responses to the oxygenic conditions varied with hypoxia type (significant *hypoxia type×clone×oxic regime* interaction in Table 3). In general, the Cold clone had the largest nuclei, followed by the Int clone, and those of the Warm clone were the smallest. The response to hypoxia was clone-specific; a decrease in cell size under hypoxic conditions was shown by Cold clone (and less apparently by the Warm clone) in the MH stage and by Int and Warm clones in the SHT stage (Fig. 3A).

**Table 3.** Final results of the GLM model for the nuclei size (a proxy for cell size) in two *L. inermis* clones.

| Effect | F-value (df num, df den) | P-value |
| --- | --- | --- |
| hypoxia type | (2, 126) 4.39 | 0.0144 |
| clone | (2, 126) 27.04 | < 0.0001 |
| oxic regime | (1, 126) 1.98 | 0.1615 |
| h. type × clone | (2, 126) 7.98 | 0.0005 |
| h. type × oxic regime | (2, 126) 7.03 | 0.0013 |
| clone × oxic regime | (2, 126) 1.29 | 0.2789 |
| h. type × clone × oxic regime | (4, 126) 14.72 | < 0.0001 |

hypoxia type – subsequent experimental stages, standardized within each stage to the SD units
oxic regime – the origin of the animals (rotifers could originate from the normoxic or hypoxic experimental conditions, but their respiration was always measured under hypoxia)

**Fig. 3.**
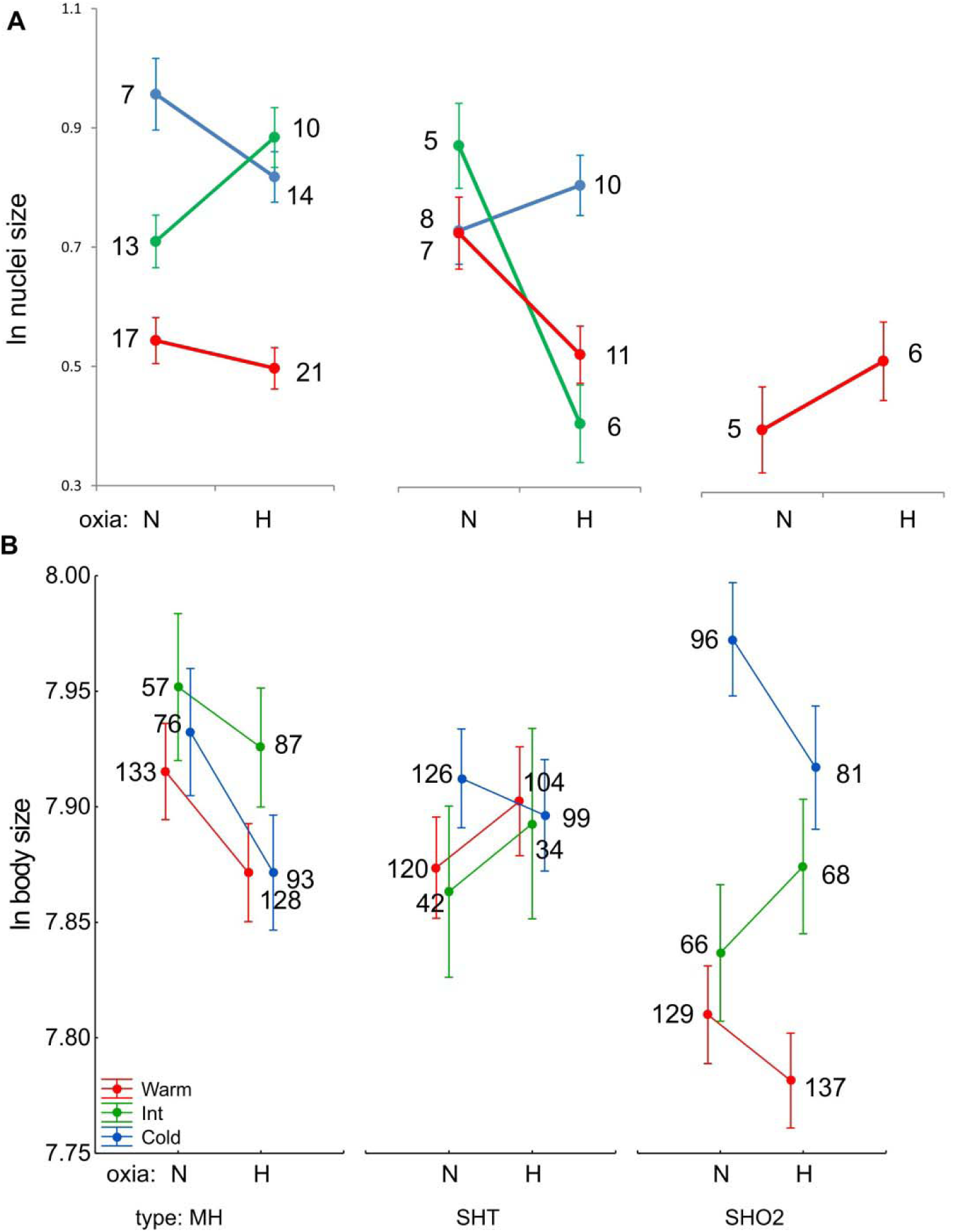
Results for cell- and body-size response. The interactions among hypoxia type (MH – mild hypoxia; SHT – severe hypoxia driven by too high of a temperature; SHO2 – severe hypoxia driven by too low of an oxygen concentration), oxic regime (N – normoxia; H – hypoxia) and clone in the GLM analysis for nuclei size (a proxy for cell size) (A), and body size (B). The numbers denote the sample size of rotifers the nuclei of which were measured (A), or the number of measured individuals (B). Least Square means ± 0.95 c.i.

### Body size

In total, we measured 1676 individuals. The mean body size of 2679 µm^2^ ± 354 SD was similar to the mean of 2411 µm^2^ shown for a different *L. inermis* clone across four thermal regimes (Walczyńska et al., 2015a). We did not find a difference between the experimental series, which was expected because of the preceding standardization. The body size was affected by the oxic regime and differed across clones. There was also a significant interaction between these two effects, and they also interacted significantly with the hypoxia type (Table 4; Fig. 3B).

**Table 4.** Final results of the GLM model for body size standardized to SD units among the hypoxia types.

| Effect | F-value (df num, df den) | P-value |
| --- | --- | --- |
| hypoxia type | (2, 1662) 1.09 | 0.3370 |
| clone | (2, 1662) 30.23 | < 0.0001 |
| oxic regime | (1, 1662) 4.76 | 0.0292 |
| h. type × clone | (4, 1662) 18.26 | < 0.0001 |
| h. type × oxic regime | (2, 1662) 9.00 | 0.0001 |
| clone × oxic regime | (2, 1662) 6.37 | 0.0018 |

hypoxia type – subsequent experimental stages, standardized within each stage to the SD units oxic regime – the origin of the animals (rotifers could originate from the normoxic or hypoxic experimental conditions, but their respiration was always measured under hypoxia)

### Fitness measures

The mean population growth rate *r* was 0.64 ± 0.08 SD indiv/day, which was similar to the value of 0.5 indiv/day that was previously shown at 30°C for a different *L. inermis* clone (Walczyńska et al., 2016). The mean egg ratio was estimated to be 0.16 ± 0.02 eggs/female. Both measures of fitness showed similar patterns across experimental regimes. The fitness differed (or tended to differ, in the case of egg ratio) across hypoxia types and clones (Table 5), with the lowest values in SHT stage and with the Warm clone showing generally the highest fitness (Fig. 4). There was a strong lack of difference between rotifers from normoxia or hypoxia in any measure (Table 5).

**Table 5.** Final results of the GLM ANCOVA models for the two fitness measures, population growth rate *r* and egg ratio.

| Effect | F-value (df num, df den) | P-value |
| --- | --- | --- |
| *population growth rate $r$: rotifers number as a dependent variable* |  |  |
| **hypoxia type** | (2, 65) 9.13 | 0.0003 |
| **clone** | (2, 65) 12.98 | < 0.0001 |
| **oxic regime** | (1, 65) 0.62 | 0.4340 |
| **initial number** | (1, 65) 11.30 | 0.0013 |
| *egg ratio: eggs number as a dependent variable* |  |  |
| **hypoxia type** | (2, 65) 3.12 | 0.0507 |
| **clone** | (2, 65) 2.99 | 0.0572 |
| **oxic regime** | (1, 65) 0.20 | 0.6587 |
| **final number** | (1, 65) 50.81 | < 0.0001 |

hypoxia type – subsequent experimental stages, standardized within each stage to the SD units oxic regime – the origin of the animals (rotifers could originate from the normoxic or hypoxic experimental conditions, but their respiration was always measured under hypoxia) the initial and final numbers – the rotifers counted in count I and count II, respectively

**Fig. 4.**
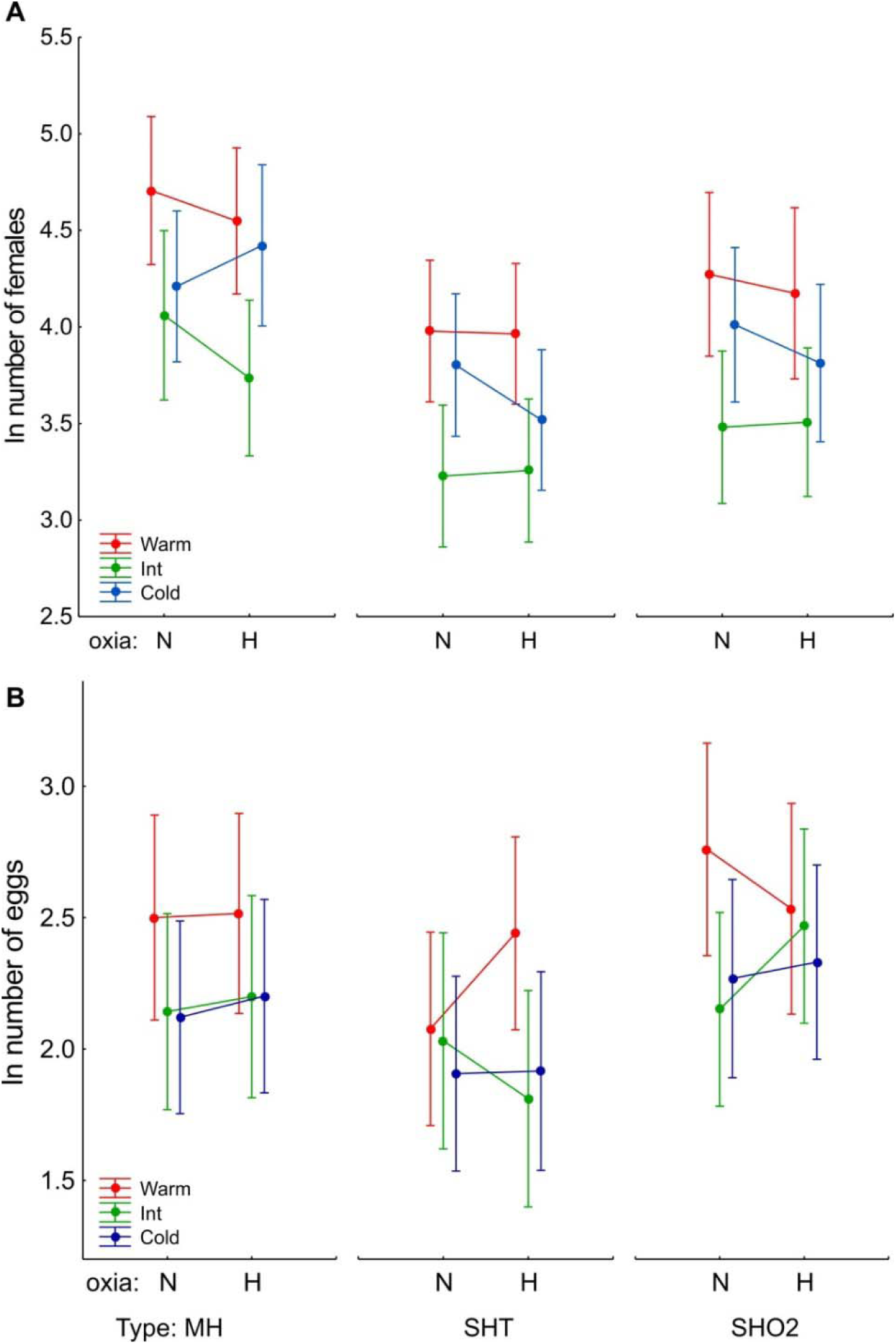
Results for the fitness measures. The interactions among the hypoxia types (MH – mild hypoxia; SHT – severe hypoxia driven by too high temperature; SHO2 – severe hypoxia driven by inadequately low oxygen concentrations), oxic regime (N – normoxia; H – hypoxia) and clone in the ANCOVA analyses for two fitness measures: population growth rate *r*, with count II number as a dependent variable and count I as a covariate (A), and egg ratio, with egg number as a dependent variable and count II as a covariate (B). N = 4 replicates. Least Square means ± 0.95 c.i.

## Discussion

The most important result from this study was that we generally confirmed the hypotheses that (i) body size decreases in response to hypoxia, (ii) as a consequence of cell size decrease, (iii) preventing the narrowing of aerobic metabolism and (iv) the subsequent decrease in fitness. Strikingly, we observed this causative pattern only for the mildly hypoxic conditions and not for the two severe hypoxia treatments (Fig. 5). In the latter case, the pattern was more complicated and clone-specific. However, in each case, the rotifers originating from the hypoxic conditions displayed a similar level of fitness to those from the normoxic conditions (Fig. 4), which could indicate some alternative mechanisms to the plastic size response that effectively prevented the decrease in fitness under stress-inducing hypoxic conditions. In this study, we confirmed the importance of the ‘optimal thermal range’ within which the TSR operates (Atkinson et al., 2003; Walczyńska et al., 2016), but we extended this issue by relating the optimality of the conditions to temperature acting in close tandem with oxygen. Additionally, the patterns we obtained concurred with a meta-analysis on 52 species of aquatic invertebrates provided by Galic et al. (2019), who found that (i) the lower the oxygen concentration was, the larger the variation in the measured response traits and that (ii) the respiration was more sensitive than growth to hypoxic conditions. We will now present the results in greater detail.

**Fig. 5.**
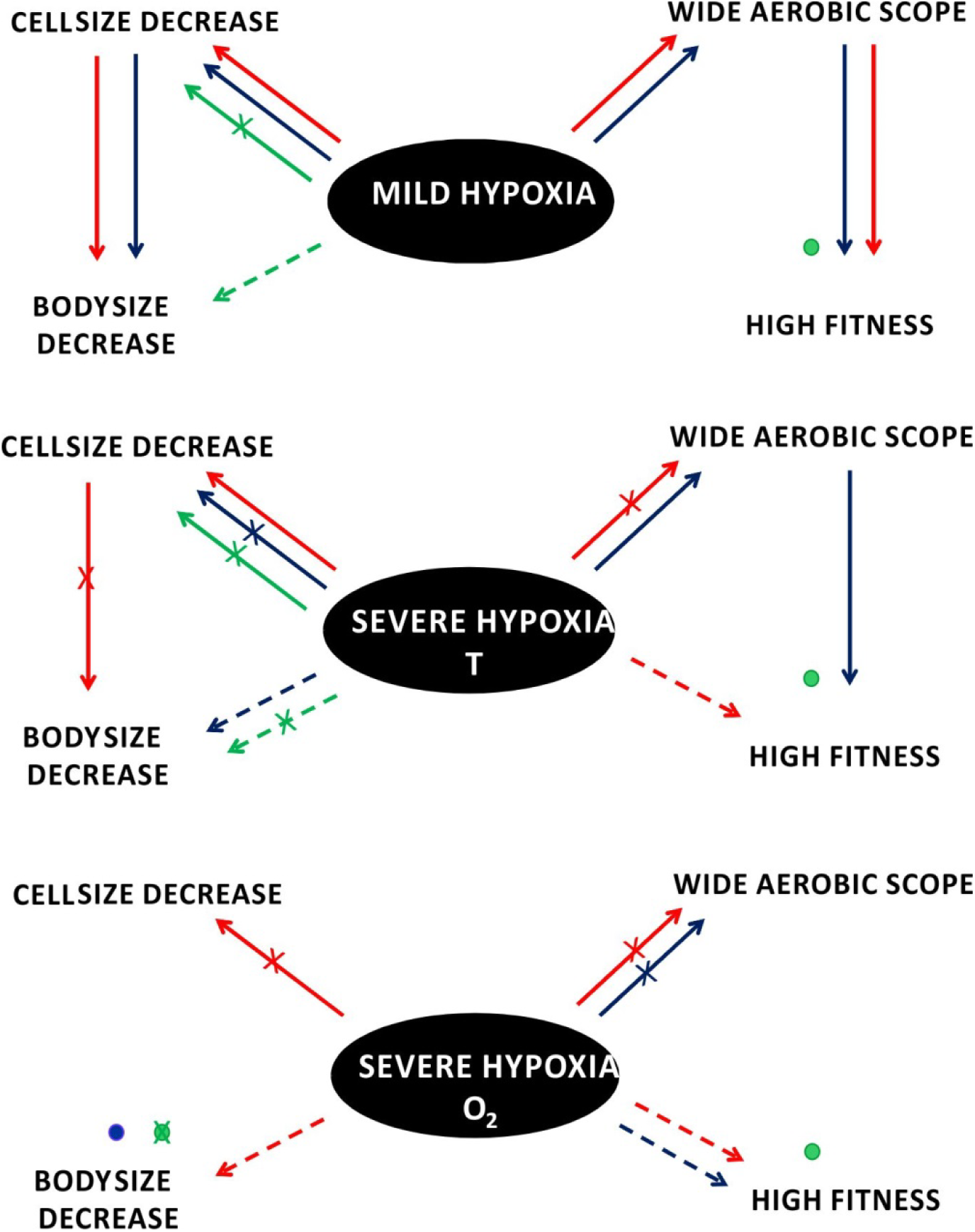
Schematic summary of the patterns found as referred to the hypotheses displayed in Fig. 1A. The arrows (red – warm-preferring clone, green – medium temperature-preferring clone, blue – cold-preferring clone) represent the positive or (when crossed) negative causative effects we found while testing across the three experimental stages. Dashed arrows indicate the effect that we found without the intermediate impact we expected. The lack of specific arrow indicates that no result was achieved for a given relationship. The dots show the positive or (when crossed) negative ultimate results in the cases when the causative effect through the intermediate trait was not known.

### Proximate mechanisms

Because of the methodological problems, we did not manage to collect the complete dataset for the cell size response. Our partial results show that in the MH treatment, the consequent body size decrease in hypoxia compared to normoxia was associated with a mechanism other than cell shrinkage in the case of one clone. In contrast, the cell and body size differences between normoxic and hypoxic rotifers showed totally opposite patterns in all clones exposed to severe temperature-driven hypoxia. A similar opposing pattern was also found in the case of Warm clone in the SHO2 treatment. The question of whether the variation in body size matches the variation in cell size was classified as fundamental by Leinaas et al. (2016). The suggestions on the differing patterns of this relationship under optimal *vs*. stressful conditions were previously invoked by Walczyńska et al. (2015a) and Walczynska et al. (2018), while the canalization *vs*. plasticity in body *vs*. cell size, which differed across *D. melanogaster*’s organs, was shown by McDonald et al. (2018). In a study in which different life stages of *D. melanogaster* were exposed to hypoxia, the cell size response was found to be stage-specific (Heinrich et al., 2011). All these results showed that the issue of body and cell size adjustment was not trivial and much can be performed in this field. The novel contribution of this study to that issue was to show that the body-cell size match was not only due to the condition but was also clone-dependent and apparently varied with thermal preferences; in a warm-preferring clone, too high temperature in the SHT treatment caused a decrease in cell size under hypoxia, but this was not followed by an adequate body size adjustment, while in the cold-preferring clone, the response was reversed under the same conditions. The intermediate clone showed a similar pattern to the warm clone in this treatment, but its body size response was opposite (to both remaining clones) in the SHO2 treatment. Based on these results, we speculated that the specific cue of plastic response to temperature-dependent hypoxia varied with organismal thermal preferences.

### Ultimate mechanisms

We found that under mild hypoxic conditions (MH treatment), the rotifers were able to adjust their metabolisms to the oxic conditions in a relatively quick time of three days (approximately 2 generations), so that the aerobic scope of the rotifers from hypoxic conditions was on average 30% wider than those of the rotifers from normoxic conditions that were exposed to acute hypoxia. The rotifers experiencing stressful hypoxic conditions for three days, either those driven by too high of a temperature (the SHT treatment) or those driven by too low of an oxygen concentration (SHO2 treatment), displayed similar (Cold clone) or even narrower (Warm clone) aerobic performances than the hypoxia-naïve rotifers (Fig. 2B, C).

Despite that, all clones performed similarly well (a fitness measure) under normoxia and hypoxia across the three hypoxia regimes (Fig. 4), though the highest average population growth rate was under mild hypoxia (approximately 15% higher than in the SHT treatment and approximately 4% higher than in the SHO2 treatment), followed by the result for the SHO2 treatment (approximately 1% higher than in the SHT treatment). The decoupling between growth (though measured at the individual level) and SDA was previously found by Chopelet et al. (2008) in *Daphnia magna* clones originating from thermally distinct geographical regions. The authors concluded that individual growth was not dictated by aerobic capacity measured as SDA. In our study, the generally worse performance displayed under severe hypoxia driven by too high of a temperature than under that driven by too low of an oxygen concentration indicated that the cue for sensing hypoxia was not indifferent for the organisms. In the case of animals that were tolerant with respect to the oxygenic conditions (Galkovskaya, 1995), the overly elevated temperatures seemed to be more stressful than too little oxygen. Our rotifers that were exposed to severe hypoxia apparently switched to alternative physiological mechanisms that we did not examine. However, it was known from previous research that this group of animals was able to switch from the regular low-efficiency and rapid lactate metabolic pathway under hypoxia to a more efficient but slower glucose-succinate metabolic pathway when exposed to severely hypoxic conditions (Esparcia et al., 1992).

### Interclonal differences

We summarize the patterns of the different strategies at the interclonal level in Fig. 5:

- Under mild hypoxia, all three clones decreased in body size, and only the Int clone used a different mechanism than cell shrinking. Regardless of the mechanism, all three clones maintained relatively high fitness under normoxia and hypoxia, and in the known cases, this was linked with keeping a wide scope for aerobic metabolism.
- Under severe hypoxia driven by too high of a temperature, each clone showed a different pattern of either decreasing cell size or body size, or, in the case of the Int clone, neither. All three clones maintained relatively high fitness despite the complex and variable body and cell size response. There was also an intriguing difference between the Cold and Warm clones: the former maintained high fitness under hypoxia by widening the aerobic scope but achieved only through body size decrease, without an adequate cell size response, while the latter displayed high performance without the widening the aerobic scope or decrease in body size under hypoxia, but with the clear pattern of cell size response.
- Under severe hypoxia driven by inadequately low oxygen concentrations, two clones decreased in body size, and in the case of the Warm clone, this response was realized without cell shrinking. None of the two clones investigated managed to maintain the wide aerobic scope under hypoxia compared to normoxia, but all three clones maintained relatively high fitness in both oxic regimes.

The reliability of our results was confirmed by the pattern found for the Warm clone, the smallest one, which displayed the highest fitness in the hot experimental conditions. An interesting image resulted from the comparison between the two measures of fitness that we applied. It seemed that in our short study, the clones exposed to stressful conditions invested either in higher fecundity (high egg ratio) or in accelerating development (high population growth rate). In the longer time scale, the two mechanisms would probably cause similar results in the overall fitness. The clearest pattern was shown by the Int clone, which had the slowest development (so that we even did not manage to examine its SDA), but its fecundity was comparable with the Cold clone across the regimes. The differences in reproductive strategy in response to hypoxia were previously found across four rotifer species by Kirk et al. (1999).

### The role of oxygen deficiency in the TSR

We concur with the conclusions from previous reports on the diverse response of ectotherms to high temperature combined with hypoxia. However, we also showed room for TSR in this general perspective; plasticity in size (decrease in high temperature/hypoxia) occurs within the relatively mild conditions, most likely as the primary mechanism of maintenance of the wide scope for aerobic metabolism. Above a certain threshold, which was previously designated by Walczyńska et al. (2016), the response changes; the plastic response was replaced by physiological mechanisms preventing, or counteracting, any possible physiological damage. Along with the growing number of empirical studies confirming the role of oxygen behind the size-to-temperature response, there has also been some criticism. These doubts are associated with the ambiguous results for higher animals, which have mostly been fishes (reviewed in Audzijonyte et al., 2019). In the study conducted on *Daphnia magna* Kielland et al. (2019) confirmed the limiting role of oxygen availability for ectotherms performance at high temperatures but concluded that plasticity did not counteract the negative effects of temperature-dependent hypoxia at very high temperatures. In our opinion, the results we present show that oxygen is the limiting factor (sensu Fry, 1947), inducing the decrease in cell size and the consequent body size decrease, but this causative relationship should be expected only under the optimal conditions, which are dependent on many factors, especially organismal life strategy and thermal preferences. Let us provide an example from everyday life. Imagine that one wants to buy a house. If there are enough funds for it, one simply buys the house; if there are not, then there are different potential strategies for gaining the necessary money: taking an additional job, saving more from the current salary, borrowing from family or friends, or taking a bank loan. In each case, the result is the same: there is enough money to buy the house. However, there are always various strategy-dependent costs, which are in each case higher than if there had been adequate funds at the beginning. The lowest-cost strategy for struggling with mild, ecologically relevant hypoxia is a plastic response. The possible and differing physiological costs for organisms struggling with severe oxygen deficiency in their environment await discovery, although we already know that they are most likely associated with hypoxia sensing at the mitochondrial level (Sokolova, 2018; Sokolova et al., 2019). A comparison of physiological responses to mild or severe hypoxia in insects was reviewed by Harrison et al. (2018). The two approaches, ecological and physiological, should be matched to determine how costly the response to temperature-dependent oxygen concentrations is, with a distinction between mild and severe conditions.

## Acknowledgments

We are very grateful to Witold Strojny for conducting the spectrophotometric analyses. The manuscript was edited by American Journal Experts. This work was supported by the National Science Centre Poland (OPUS 2015/19/B/NZ8/01948) and by the Jagiellonian University (DS/INoS/757/2019). **The authors declare no conflict of interest.**

## Data availability

The data supporting the results will be archived in a public repository of the Jagiellonian Univeristy and the data DOI will be included at the end of the article. The data will be made publicly available at the time of publication.

---

**Fig. S1.**
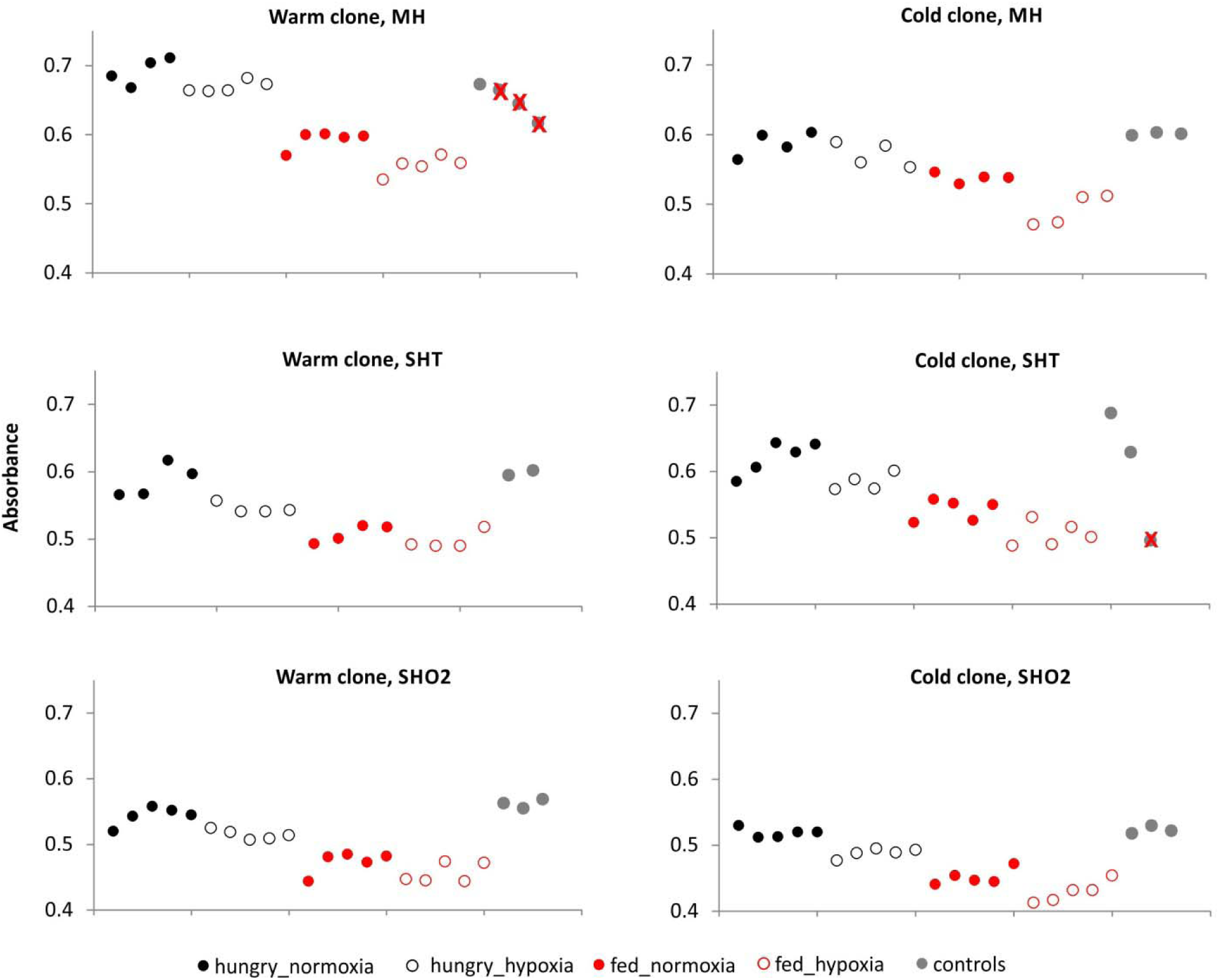
The analysis of raw absorbance data for the reliability of controls. The control samples perceived as unreliable are crossed out and were not included in the analyses. Absorbance is a measure proportional to the amount of oxygen dissolved in a sample. MH – mild hypoxia treatment, SHT – severe hypoxia treatment, driven by elevated temperature, SHO2 – severe hypoxia treatment driven by alleviated oxygen concentration.

